# Evolution of color phenotypes in two distantly related species of stick insect: different ecological regimes acting on similar genetic architectures

**DOI:** 10.1101/023481

**Authors:** Aaron A. Comeault, Clarissa Ferreira, Stuart Dennis, Víctor Soria-Carrasco, Patrik Nosil

## Abstract

Recurrent (e.g. parallel or convergent) evolution is widely cited as evidence for natural selection’s central role in evolution but can also highlight constraints affecting evolution. Here we describe the evolution of green and melanistic color phenotypes in two species of stick insect: *Timema podura* and *T. cristinae*. We show that similar color phenotypes of these species (1) cluster in phenotypic space and (2) confer crypsis on different plant microhabitats. We then use genome-wide association mapping to determine the genetic architecture of color in *T. podura*, and compare this to previous results in *T. cristinae*. In both species, color is under simple genetic control, dominance relationships of melanistic and green alleles are the same, and SNPs associated with color phenotypes colocalize to the same genomic region. These results differ from those of ‘typical’ parallel phenotypes because the form of selection acting on color differs between species: a balance of multiple sources of selection acting within host species maintains the color polymorphism in *T. cristinae* whereas *T. podura* color phenotypes are under divergent selection between hosts. Our results highlight how different adaptive landscapes can result in the evolution of similar phenotypic variation, and suggest the same genomic region is involved.

## Introduction

Recurrent evolution, such as parallel or convergent evolution, occurs when functionally or genetically similar phenotypic variation evolves repeatedly in multiple populations or species (Colosimo *et al.* 2005; Arendt & Reznick 2008; Deagle *et al.* 2012; Pfenninger *et al.* 2014). Recurrent evolution of phenotypic variation that overlaps in phenotypic space, under similar ecological conditions, has been widely used to argue for natural selection’s role as a fundamental driver of evolution (Schluter 2000; Losos 2011). However, similar phenotypes might also repeatedly evolve under different ecological conditions when genetic or developmental biases exist that constrain the range of possible phenotypic outcomes (Gould & Vrba 1982; Smith *et al.* 1985; Arnold 1992; Price & Burley 1994; Schluter 1996). Thus, a better understanding of repeated evolution in response to both similar and different ecological contexts can shed light on constraints in evolution. Classically, studies of recurrent evolution have focused on convergent or parallel phenotypic traits that have evolved in response to similar selective regimes (e.g. Colosimo *et al.* 2005; Castoe *et al.* 2009; Quek *et al.* 2010; Losos 2011; Reed *et al.* 2011; Foote *et al.* 2015). By contrast, we generally lack examples of phenotypes that have repeatedly evolved in response to different adaptive landscapes.

In addition to ecological processes, the genetic mechanisms underlying recurrent evolution can involve different sources of genetic variation (Rosenblum *et al.* 2010, 2014; Manceau *et al.* 2010). As a first source, recurrent evolution can be due to independent mutations and these may arise at the same site in the genome, at different sites within the same gene, or within different genes in the same genetic network (Karasov *et al.* 2010; Manceau *et al.* 2010). Second, recurrent evolution can be derived from the repeated ‘quasi-independent’ sorting of pre-existing alleles segregating as standing genetic variation (Barrett & Schluter 2008). For example, both threespine stickleback (Schluter & Conte 2009) and *Heliconius* butterflies (Dasmahapatra *et al.* 2012; Martin *et al.* 2013) have repeatedly evolved similar phenotypes through selection acting on standing variation. The distinction between these genetic mechanisms is also of theoretical interest because evolution from standing variation can be more rapid, particularly in smaller, mutation-limited populations (Barrett & Schluter 2008; Schluter & Conte 2009; Barton 2010; Karasov *et al.* 2010; Uecker & Hermisson 2011). In this context, quantitatively describing the genetic architecture of phenotypes involved in recurrent evolution is a first step towards resolving these mechanisms (Arendt & Reznick 2008).

Here we study the ecological and genetic mechanisms of recurrent evolution using two species of *Timema* stick insects. The genus *Timema* is comprised of ∼21 species of herbivorous insects that are endemic to southwestern North America and show a wide range of within- and among-species variation in body coloration (Sandoval *et al.* 1998). Frequently, the same colors are present in distantly related species of *Timema* (Crespi & Sandoval 2000), providing a well suited system for addressing questions about the ecological and genetic basis of recurrent evolution. Here we focus on a green / melanistic color polymorphism that is found within two distantly related species with completely non-overlapping geographic ranges: *T. cristinae* and *T. podura* (Fig. 1). These species are estimated to have diverged from a common ancestor approximately 20 million years ago (*Timema* have a single generation per year; Sandoval *et al.* 1998). Therefore, they represent an interesting system in which to study recurrent evolution because it is unclear if a shared genetic basis is expected between species that diverged so long ago (Conte *et al.* 2012). Details of the two species are as follows.

**Figure 1.**
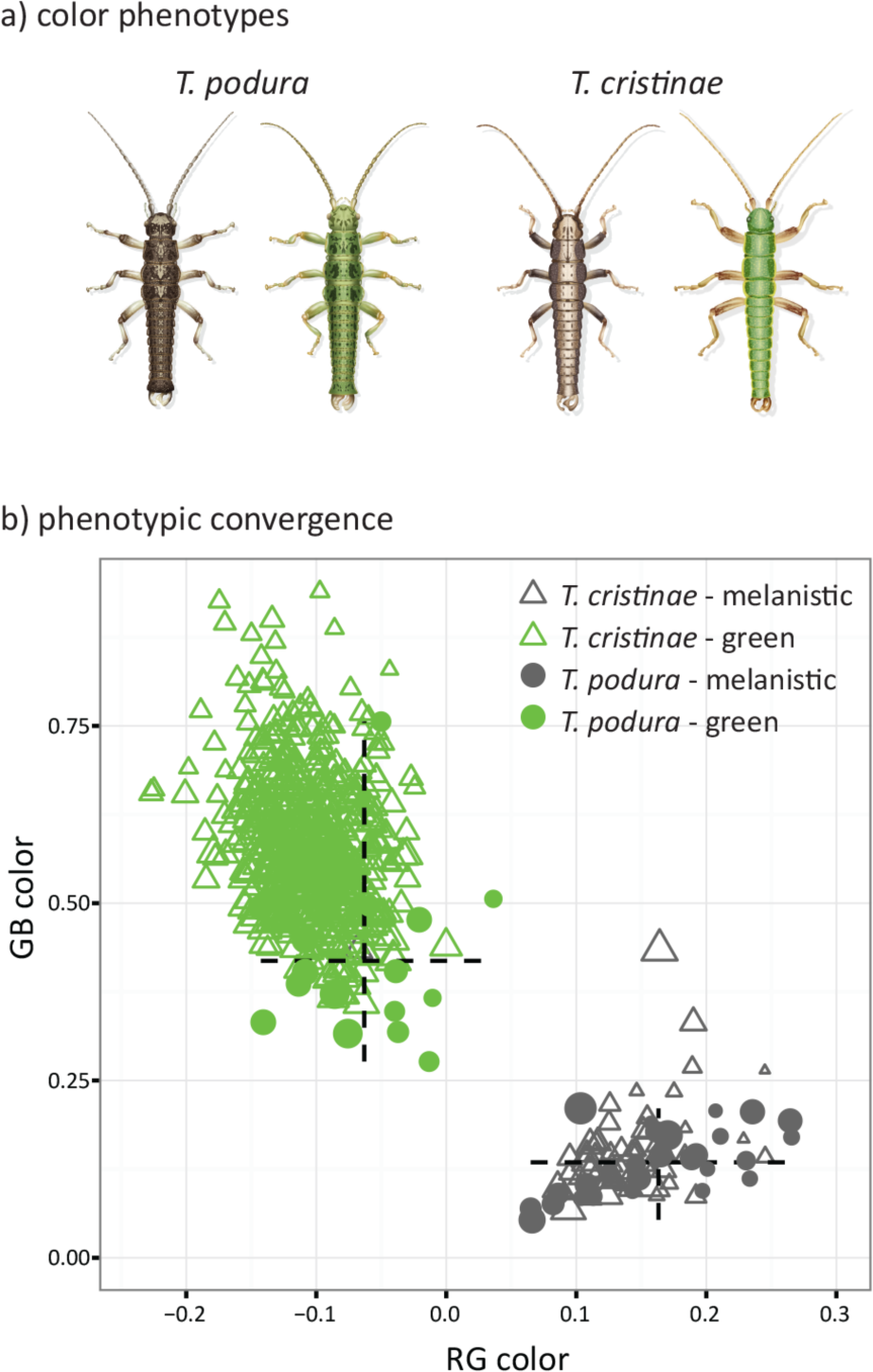
(a) Illustrations of melanistic and green phenotypes for *T. podura* and *T. cristinae* (illustrations are of males; credit: Rosa Marin). (b) Phenotypic position of 42 *T. podura* and 602 *T. cristinae* in RG – GB color space. Hashed lines in ‘b’ represent the range of RG (horizontal line) and GB (vertical line) values for *T. podura* phenotypes and size of the symbols is proportional to an individual’s luminance.

*Timema cristinae* is endemic to the coastal chaparral of the westernmost mountains of the Transverse Ranges of southern California and is found on the two primary host plants (*Ceanothus spinosus* and *Adenostoma fasciculatum*). Within populations found on both host species, green and melanistic color phenotypes segregate as a polymorphism and the frequency of the color phenotypes does not differ between host species (Comeault *et al.* 2015). Evidence indicates that these color phenotypes are maintained by a balance of selective agents that are similar between hosts and include selection for crypsis in leafy (green favored) and woody (melanistic favored) plant microhabitats, differences in fungal infection between color phenotypes, and potential fitness differences associated with climatic variation. Segregation of color in classical genetic crosses and genome wide association (GWA) mapping indicate that *T. cristinae* color phenotypes are controlled by a simple genetic architecture with most variation explained by a single genetic region localized on a single linkage group (LG8), with the green allele dominant to the melanistic allele (Comeault *et al.* 2014, 2015).

The other species we consider, *T. podura*, is endemic to the San Bernardino, Rosa and San Jacinto Mountains of central southern California and also inhabits host plant species in the genus *Ceanothus* (*C. leucodermis*) and *Adenostoma* (*A. fasciculatum*). Like *T. cristinae*, *T. podura* has both a green and a melanistic color morph (Fig. 1; melanistic individuals have also been referred to as “grey” or “red”; Sandoval & Nosil, 2005). However, unlike *T. cristinae* the frequency of *T. podura* color phenotypes is different between populations living on different host species: green *T. podura* are, to our knowledge, not found on *A. fasciculatum* (Sandoval & Nosil 2005). Moreover, experiments have shown that avian predators preferentially depredate melanistic individuals when on *C. leucodermis* (potentially due to the light green color of *C. leucodermis* branches, an hypothesis we test here) and green individuals on *Adenostoma* (Sandoval & Nosil 2005). Thus, in contrast to *T. cristinae*, there is evidence for divergent selection acting on *T. podura* color phenotypes between host species. The maintenance of melanistic *T. podura* on *C. leucodermis* could be due to gene flow between populations found on different hosts, as documented in *T. cristinae* at spatial scales similar to those separating populations of *T. podura* on different hosts (Nosil *et al.* 2012; Sandoval and Nosil 2005). In *T. podura*, quantitative tests for crypsis of different color phenotypes between different host species and plant microhabitats are lacking and the genetic basis of color phenotypes is unknown.

Here we present analyses of color variation within a polymorphic population of *T. podura* living on the host plant *C. leucodermis*. We first quantify color phenotypes in both *T. podura* and *T. cristinae* to test whether they overlap in phenotypic space. We then determine host plant microhabitats in which the color phenotypes confer the greatest degree of crypsis. In addition to phenotypic analyses, we conducted multi-locus GWA mapping to quantitatively describe aspects of the genetic architecture of *T. podura* color phenotypes, such as whether they are controlled by few or many loci and the nature of any dominance relationships between alleles. Finally, we map single nucleotide polymorphisms (SNPs) associated with color variation in *T. podura* to the *T. cristinae* reference genome, testing if they co-localize to the same genomic region as SNPs associated with color in *T. cristinae*. Finally, we describe the genes found on scaffolds containing SNPs associated with color variation. Our results inform the ecological and genetic basis of recurrent evolution and pave the way for future functional studies characterizing genes causally involved in adaptation in the genus *Timema*.

## Methods

### Quantifying variation in color

We recorded digital images of 42 adult *T. podura* collected from a phenotypically variable population collected off of *C. leucodermis* plants (population code: BSC; latitude 33.816, longitude -116.790). Images were recorded in RAW format with a Canon 600d camera equipped with an EF-S 60mm F2.8 macro lens (Canon (UK) Ltd., Surry, UK) with an aperture of f/14, a shutter speed of 1/250s, and two wireless external flashes (Yongnuo YN560-II speedlight, Yongnuo Digital, www.yongnuo.eu) set to manual. Each image was captured with 1 cm grid paper as a background and included a standard color chip (Colorgauge Micro, Image Science Associates LLC, Williamson, NY, USA). Following image capture each individual was placed in an individually labeled 1.5 ml microcentrifuge tube and preserved in 95% ethanol for subsequent DNA extraction. Images were linearized and corrected to 80% reflectance based on a neutral gray color target (target #10 of the cologauge micro color chip) with Adobe’s Photoshop Lightroom 4 software (Adobe Systems Software Ireland Ltd), and saved as a .tif file. To quantify color in *T. cristinae* we used 602 images of *T. cristinae* that were collected using the same protocol outlined above in a previous study, but used only to qualitatively describe color in *T. cristinae* (classified as green versus non-green body coloration; Comeault *et al.* 2015). Here, we report novel analyses of these photos based on quantitative measurements of color.

In addition to images of insects, we recorded digital images of plant microhabitats frequently encountered by *Timema* in nature, which have not been analyzed in past work. Specifically, for *T. podura* we recorded images of the host plants *C. leucodermis* and *A. fasciculatum*, and for *T. cristinae* we recorded images of *C. spinonus* and *A. fasciculatum*. For each host plant we recorded images of the leaf leaves and wood. For *Ceanothus spp.*, images of both the top and bottom of the leaves were recorded. *A. fasciculatum* leaves are needle-like and do not have a definable top or bottom; therefore we did not define a leaf top or bottom for this host species. From host plants collected from the same location as the *T. podura* included in this study (33.816, -116.790) we recorded 11 images for the top and bottom of *C. leucodermis* leaves, 10 images of *C. leucodermis* wood, 11 images of *A. fasciculatum* leaves and 10 images of *A. fasciculatum* wood. From host plants collected from the same location as *T. cristinae* (34.518, -119.801) we recorded 5 and 6 images of the top and bottom of *C. spinosus* leaves, respectively, 5 images from the wood of *C. spinosus*, and 6 images of the leaves and wood of *A. fasciculatum* plants.

To quantify color we first recorded RGB values from linearized and corrected digital images. For *T. podura* and *T. cristinae* we measured color on the lateral margin of the second thoracic segment, and for plant microhabitats we measured areas of the given microhabitat that lacked shadow and glare, using the polygon selection tool in ImageJ (Abràmoff *et al.* 2004). We then recorded mean RGB values for each color patch using the color histogram plugin in ImageJ. For each color patch we converted raw RGB values to variables representing two color channels and one luminance channel as suggested by Endler (2012). A red-green color channel (RG) was calculated using the relationship (R-G)/(R+G), a green-blue color channel (GB) as (G-B)/(G+B), and a luminance (i.e. brightness; L) channel as (R+G+B) for each individual and microhabitat type (Endler 2012). While this method of measuring color does not take into account the visual system of the receiver or the light environment an object is viewed in, it does represent an unbiased quantification of color that can be useful in a comparative context. Moreover, *T. cristinae* does not reflect UV light (Comeault *et al.* 2015) indicating that the digital images we use here likely capture a majority of the biologically relevant differences between the color phenotypes.

Using RG, GB, and L values we quantified phenotypic overlap in color between *T. podura* and *T. cristinae*. First, we used linear models to compare RG, GB, and L values between green and melanistic *T. podura* color phenotypes, green and melanistic *T. cristinae* color phenotypes, green *T. podura* and green *T. cristinae*, and melanistic *T. podura* and melanistic *T. cristinae*. We also analyzed the position of the different color phenotypes in phenotypic space using an approach analogous to that used by Beuttell & Losos (1999) to quantify clustering of *Anolis* ecomorphs in multivariate phenotypic space. Specifically, we first calculated the Euclidean distance between all individuals in our sample (i.e. all pairwise comparisons) in RG – GB color space. We then used Wilcoxon signed rank tests to determine differences in color distance between the same colored individuals of the two species (i.e., the distances between green *T. podura* and green *T. cristinae* or melanistic *T. podura* and melanistic *T. cristinae*) and either different colored individuals of *T. cristinae* or different colored individuals of *T. podura*. These analyses enabled us to ask whether the same color phenotypes of the two species are closer to each other, in phenotypic space, than to the alternate color phenotype of their own species.

To estimate the degree of color matching between the two color morphs of *T. podura* and different plant microhabitats we calculated three measures of ‘color matching’ by subtracting individual *T. podura* color (RG and GB) and luminance (L) values from those of each of the five different plant microhabitats. We then determined whether the different color phenotypes of *T. podura* differed in their degree of color matching to different host plant microhabitats by comparing the difference between RG, GB, and L values of different individual / plant-microhabitat combinations using nested analysis of variance (ANOVA). We have previously described levels of color matching between color phenotypes of *T. cristinae* and their host plant microhabitats using reflectance spectra (Comeault *et al.* 2015). However, we here carried out a parallel analysis in *T. cristinae* based on the measures from photographs as described above. This allowed us to confirm results obtained from reflectance spectra and to directly compare results to those from *T. podura*. Within *T. cristinae*, we recovered qualitatively similar results when using the RG, GB, and L metrics to those obtained from color analyses of reflectance spectra that take into account both the visual system of the receiver and the ambient light environment (see Fig. S1 and Comeault *et al.* 2015). All statistical analyses were carried out in R (Core Team 2013).

### Genomic sampling

We extracted whole genomic DNA from 50 *T. podura* (19 green and 31 melanistic) that included the same 42 individuals used to quantify color from photographs and 8 additional individuals sampled from the same population (qualitatively scored as “green” or “melanistic”) using Qiagen DNeasy blood and tissue kits (Qiagen). Following the method of Parchman et al. (2012), which we have applied to *Timema* in the past (Nosil *et al.* 2012; Comeault *et al.* 2014; Gompert *et al.* 2014), we created individually barcoded restriction-site associated DNA libraries for sequencing on the Illumina HiSeq platform. Briefly, these libraries are generated by digesting genomic DNA in the presence of the restriction enzymes *Eco*RI and *Mse*I (New England Biolabs), ligating double stranded adapters containing the Illumina priming site and one of 50 unique 8 to 10 base pair (bp) barcode sequence to the restriction fragments, and amplifying ligated fragments using the polymerase chain reaction (PCR). For a detailed protocol refer to Parchman et al. (2012). We then pooled these 50 libraries with an additional 48 uniquely barcoded libraries that were part of another study. The pooled libraries were selected for fragments ranging in size from 300 to 500 bp with Pippin-prep targeted size selection (Sage Science, Inc., MA, USA) and sequenced on a single lane of the Illumina HiSeq2000 platform using V3 reagents at the National Center for Genome Research (Santa Fe, NM, USA). Sequenced libraries were used to generate a genotype-by-sequencing (GBS) dataset that allowed us to map the genetic basis of *T. podura* color phenotypes.

We removed barcodes and the following six bp of the *Eco*RI cut site from raw sequence reads, while allowing for single bp errors in the barcode sequence due to synthesis or sequencing error, using a custom Perl script developed and implemented in Nosil *et al.* (2012). Following removal of barcode sequences this resulted in a total of 130,280,785 raw sequence reads with an average of 2,605,616 reads per individual (95% interval = 1,351,013 – 3,356,050) and an average length of 83 bp (95% interval = 63 – 86). We aligned 90,923,479 of these reads (69.8%) to the reference genome sequence of *T. cristinae* (Soria-Carrasco *et al.* 2014) using bowtie2 version 2.1.0 (Langmead & Salzberg 2012) with the local model and the ‘--very-sensitive-local’ preset (-D 20 -R 3 -N 0 -L 20 -i S,1,0.50). We used samtools version 0.1.19 (Li *et al.* 2009) to sort and index alignments. We used the reads mapped to the *T. cristinae* genome to generate a reference consensus sequence of *T. podura* using samtools mpilup and bcftools. We used vcfutils.pl with the vcf2fq command to filter out positions with a number of reads below 8 and above 500, as well as those with a phred-scale mapping quality score lower than 20. Filtered sites were coded as missing data. Subsequently, we used bowtie2 with the same arguments used above to align 100,095,223 raw reads (76.8%) to this reference consensus. As before, the alignments were sorted and indexed with samtools.

Variants were called using samtools, mpileup, and bcftools using the full prior and requiring the probability of the data to be less than 0.5 under the null hypothesis that all samples were homozygous for the reference allele to call a variant. Insertion and deletion polymorphisms were discarded. We identified 638,828 single nucleotide polymorphisms (SNPs) that were reduced to 137,650 SNPs after discarding SNPs for which there were sequence data for less than 40% of the individuals, low confidence calls with a phred-scale quality score lower than 20, and SNPs with more than two alleles. Average depth of the retained SNPs across all individuals was ∼460x (mean coverage per SNP per individual ∼ 9x). We used a custom Perl script to calculate empirical Bayesian posterior probabilities for the genotypes of each individual and locus using the genotype likelihoods and allele frequencies estimated by bcftools along with Hardy-Weinberg priors (i.e. p(A)=p^2^; p(a)=(1-p)^2^; p(Aa)= 2p(1-p)). Finally, we computed the posterior mean genotype scores for each individual, at each locus, by multiplying the probability of the homozygous minor allele genotype by two and adding the probability of the heterozygous genotype. These imputed genotype scores range from zero to two and represent the dosage of the minor allele in a given genotype. All imputed genotype scores were saved in bimbam file format and used for GWA mapping analyses. Principal component analysis based on imputed genotype scores for these 50 individuals revealed a lack of genetic structure (see Fig. S2 of SI).

### *Genetic architecture of* T. podura *color phenotypes estimated using GWA*

We estimated aspects of the genetic architecture of color variation in *T. podura* using multi-locus Bayesian sparse linear mixed models (BSLMMs) as implemented in the software package *gemma* (Zhou & Stephens 2012; Zhou *et al.* 2013). Because *T. podura* color phenotypes were completely non-overlapping in two-dimensional color space (Fig 1b) we unambiguously scored each of the 50 genotyped individuals as green (*n* = 19) or melanistic (*n* = 31) and ran probit BSLMMs in *gemma* (as done for green and melanistic phenotypes of *T. cristinae* in Comeault *et al.* 2015). BSLMMs allow for multi-SNP mapping and can be used to estimate three hyperparameters that describe aspects of the genetic architecture of a given trait (Zhou & Stephens 2012; Zhou *et al.* 2013). First, the model estimates the total phenotypic variation explained by all the SNPs included in an analysis (proportion of phenotypic variation explained; PVE). PVE is therefore an estimate of the combined effect of SNPs with both ‘large’ (i.e., detectable) phenotypic effects and ‘polygenic’ (i.e., infinitesimal and undetectable) SNPs with minor effects on phenotypic variation. Second, *gemma* estimates the proportion of the total phenotypic variation (i.e. PVE) that can be explained by ‘large-effect’ SNPs alone (proportion of genetically-explained variation; PGE). Third, *gemma* estimates the number of SNPs (n-SNP) that have non-zero effects on phenotypic variation (i.e. the number for which the relationship between genotype and phenotype [β] is greater than zero). In essence, the estimate of n-SNP represents the number of ‘large-effect’ SNPs needed to explain the PGE. We implemented BSLMMs in *gemma* with 10 independent Markov-chain Monte Carlo (MCMC) chains ran for 25 million steps with an initial burn-in period of 5 million steps. Parameter values estimated by the BSLMMs were recorded every 100 steps and written every 10,000 steps. All additional options in *gemma* remained at default values and SNPs with minor allele frequencies < 0.01 were excluded from these analyses. Here we report the median and 95% credible interval (95% equal tail posterior probability intervals [95% ETPPIs]) for PVE, PGE, PVE x PGE (an estimate of the total phenotypic variation explained by only SNPs with large phenotypic effects), and n-SNP.

We carried out analyses to test the strength of the genetic signal in our data set to accurately estimate hyperparameters. First, we conducted a permutation test using GWA mapping in *gemma* as described above with five data sets generated by randomly permuting phenotypic scores for each individual. Second, we performed cross validation using the genomic prediction function in *gemma* to predict phenotypes of individuals whose color phenotype was randomly masked from our data set (SI for details).

In addition to the hyperparameters described above, *gemma* provides the posterior inclusion probability (PIP) and estimates the phenotypic effect (β) of each SNP that is identified to have a non-zero effect on phenotypic variation in at-least one model iteration. As such, PIP is computed as the proportion of BSLMM iterations for which a given SNP is identified as having a non-zero β. Therefore, SNPs that are more strongly associated with phenotypic variation will have larger PIPs and these SNPs are the strongest candidates of being linked to the functional variant(s) underlying phenotypic variation. Here we focus on high PIP as evidence that a given SNP is associated with variation in *T. podura* color phenotypes.

### Co-localization of regions associated with color in the two species

To determine whether SNPs associated with color phenotypes (‘candidate SNPs’ hereafter) in *T. podura* map to similar genomic regions as those in *T. cristinae*, we localized genetic effects by calculating the mean PIP of SNPs at two different genomic scales. First, we calculated the mean PIP of SNPs mapping to each of the 13 *T. cristinae* linkage groups (LGs) (Soria-Carrasco *et al.* 2014). Second, we calculated the mean PIP of SNPs mapping to each of the 1413 scaffolds (1311 of which contained SNPs in our data sets) that make up the 13 *T. cristinae* LGs.

Because we found SNPs mapping to LG 8 to have the largest mean PIP in both *T. cristinae* and *T. podura* (see Results), we assessed the likelihood that this pattern would happen by chance using permutation tests. The purpose of these analyses was to determine the probability that co-localization of SNPs with high PIPs is expected by chance when we account for (1) the genomic distribution of SNPs in our data set and (2) the distribution of PIPs observed for these SNPs. To determine whether there is statistical evidence for clustering of PIPs at the level of LGs we randomly permuted PIPs without replacement 10,000 times for both the *T. podura* and *T. cristinae* SNP data sets. During this permutation procedure the number and location of SNPs along each linkage group was maintained and therefore the only difference in these 10,000 data sets was the distribution of PIPs across the genome. We calculated the proportion of permuted data sets for which LG 8 had the largest mean PIP in both species.

We also tested whether there was evidence for the co-localization of candidate SNPs within LG 8 by calculating the mean distance between two randomly sampled scaffolds along LG 8. The distance between randomly sampled scaffolds was measured in terms of the number of scaffolds separating the two sampled scaffolds because the absolute distance between scaffolds (in bp) in the current version of the *T. cristinae* genome is unknown. We repeated this procedure 10,000 times and determined the probability that the distance observed between two randomly sampled scaffolds would be less than or equal to the minimum observe distances between the top two *T. cristinae*, and the top two *T. podura* candidate scaffolds.

### Dominance relationships between alleles

We next determined dominance relationships of alleles at candidate SNPs in *T. podura* using methods previously applied to *T. cristinae* (Comeault *et al.* 2015). Specifically, we computed the ratio of dominant to additive effects of alleles at each of the loci identified by BSLMMs as having high PIPs. Dominance effects (*d*) are calculated as the difference between the mean phenotype of heterozygotes and half the phenotypic distance between the mean phenotypes of the two homozygous genotypes. Additive effects (*a*) were calculated as half the phenotypic distance of the two homozygous genotypes. The ratio *d*/*a* represents the deviance of the phenotypes of heterozygotes from those expected under additivity (Burke *et al.* 2002; Miller *et al.* 2014). The expected value of *d*/*a* for additive alleles is 0 while completely dominant or recessive alleles will be 1 or -1. Here we follow previous conventions (Burke *et al.* 2002; Miller *et al.* 2014) and classify alleles as being dominant if *d*/*a* is greater than 0.75, recessive if *d*/*a* is less than -0.75, partially dominant or partially recessive if *d*/*a* is between 0.75 and 0.25 or -0.75 and -0.25, respectively, and additive if *d*/*a* is between - 0.25 and 0.25.

LD among sequenced SNPs can influence individual PIPs and be used to assess whether SNPs associated with phenotypic variation are tagging the same functional variant or independent genetic variation. For example, consider two functionally neutral SNPs that are in perfect LD with each other and a single causal SNP that explains some amount of phenotypic variation but was not itself sequenced. Each of the two neutral SNPs could be identified by BSLMMs in ∼50% of model iterations and would therefore each have individual PIPs of 0.5. However, taken together, these two SNPs would be tagging the single causal SNP in 100% of model iterations (50% + 50%). To address whether candidate SNPs associated with color variation were potentially tagging the same causal genetic variant (or causal variants in LD with one another) we estimated LD by calculating genotypic correlations (*r*^2^) and normalized disequilibrium coefficients (D’) among different groups of SNPs using the ‘r2fast’ and ‘dprfast’ functions of the *GenABEL* library in R (Aulchenko *et al.* 2007), respectively. To determine whether levels of LD between the candidate SNPs were greater than null genomic expectations we calculated pairwise *r*^2^ and D’ among all SNPs located on candidate scaffolds 1806 and 284, 1000 SNPs randomly sampled from LG 8, and 1000 SNPs randomly sampled from the genome.

### Functional annotation of genomic regions containing candidate SNPs

To generate a list of candidate genes underlying color phenotypes in *T. podura* we examined annotations of predicted genes in the current draft of the *T. cristinae* reference genome for both *T. podura* candidate scaffolds (i.e. scaffold 284 and 1806; see Results). We then manually extracted and tabulated all InterPro or GO annotations for each predicted gene located on these two scaffolds.

Previous work has shown that a SNP located in an intron of a gene encoding a cysteinyl-tRNA synthetase is strongly associated with color phenotypes in *T. cristinae* (Comeault *et al.* 2015). The GWA mapping analyses carried out here revealed that SNPs strongly associated with color in *T. podura* (i.e. the two candidate SNPs identified on scaffolds 284 and 1806) map to different scaffolds than the scaffold containing the *T. cristinae* candidate cysteinyl-tRNA synthetase gene (scaffold 842; see Results below). We therefore determined whether any of the SNPs within the *T. podura* GBS data set are found within, or near, this candidate gene. Because the *T. podura* data set does not contain any SNPs mapping to this gene (see Results), we also determined whether the *T. podura* SNPs mapping to scaffold 842 were in strong LD with the two candidate SNPs we identify here. The goal of these analyses was to determine the likelihood that our data set contained SNPs that would recover a relationship between color and the candidate scaffold identified in *T. cristinae*, should it exist.

## Results

### Quantifying variation in color

The green and melanistic phenotypes of *T. podura* differ in both RG and GB color values (*F*_1, 40_ = 158.92, *P* < 0.001; *F*_1, 40_ = 126.66, *P* < 0.001) but not with respect to luminance (*F*_1, 40_ = 3.76, *P* = 0.06). Color phenotypes of *T. cristinae* differ in RG color, GB color, and luminance (RG color: *F*_1, 600_ = 2687.30, *P* < 0.001; GB color: *F*_1,_ 600 = 1050.90, *P* < 0.001; luminance: *F*_1,_ 600 = 52.07, *P* < 0.001). Melanistic *T. podura* do not differ from melanistic *T. cristinae* in RG or GB color (*t* = 1.88, *P* = 0.07; *t* = -1.52, *P* = 0.13) but melanistic *T. podura* have significantly greater luminance than melanistic *T. cristinae* (mean L = 240.98 and 178.62, respectively; *t* = 5.12, *P* < 0.001). Green *T. podura* differ in RG color, GB color, and L from the green phenotype of *T. cristinae*, (*t* = 3.33, *P* = 0.004; *t* = -5.75, *P* < 0.001; and *t* = 5.48, *P* < 0.001, respectively).

Despite some difference in color between *T. podura* and *T. cristinae*, both green and melanistic color phenotypes broadly overlap in RG – GB color space and the Euclidean distances between similarly colored individuals of each species were much less than the Euclidean distances between differently colored individuals within species (mean [SE] Euclidean distance between *T. podura* and *T. cristinae* having the same color = 0.193 [0.0011] and between differently colored *T. podura* = 0.377 [0.0050] or *T. cristinae* = 0.501 [0.0006]; Fig. 1b). Therefore, while there are slight differences in the color of green *T. podura* and green *T. cristinae*, these phenotypes cluster tightly in phenotypic space and are more similar in color to each other than to differently colored individuals of their own species (*U* = 315,985,777, *P* > 0.0001; Fig. 1b).

Color phenotypes of *T. podura* differed significantly in their degree of color matching against different plant microhabitats (nested ANOVA; RG color: *F*_1,40_ = 158.9, *P* < 0.001; GB color: *F*_1,40_ = 126.7, *P* < 0.001). Color phenotypes do not differ with respect to luminance when compared against different plant microhabitats (nested ANOVA; L: *F*_1,40_ = 3.8, *P* = 0.06). Figure 2 shows that the RG color of green *T. podura* matches the RG color of all leaf microhabitats and the wood of *C. leucodermis* more closely than the RG color of the melanistic phenotype. Similarly, the GB color of green *T. podura* matches the color of *A. fasciculatum* leaves and the top of *C. leucodermis* leaves more closely than the GB color of the melanistic phenotype. RG and GB color of melanistic *T. podura* closely matches RG and GB color of *A. fasciculatum* wood and the GB color of melanistic *T. podura* also matches the GB color of *C. leucodermis* wood (Figure 2). These results show that the color of melanistic *T. podura* poorly matches both *C. leucodermis* microhabitats (bark and leaves), but closely matches the color of *A. fasciculatum* bark, relative to the green phenotype.

**Figure 2.**
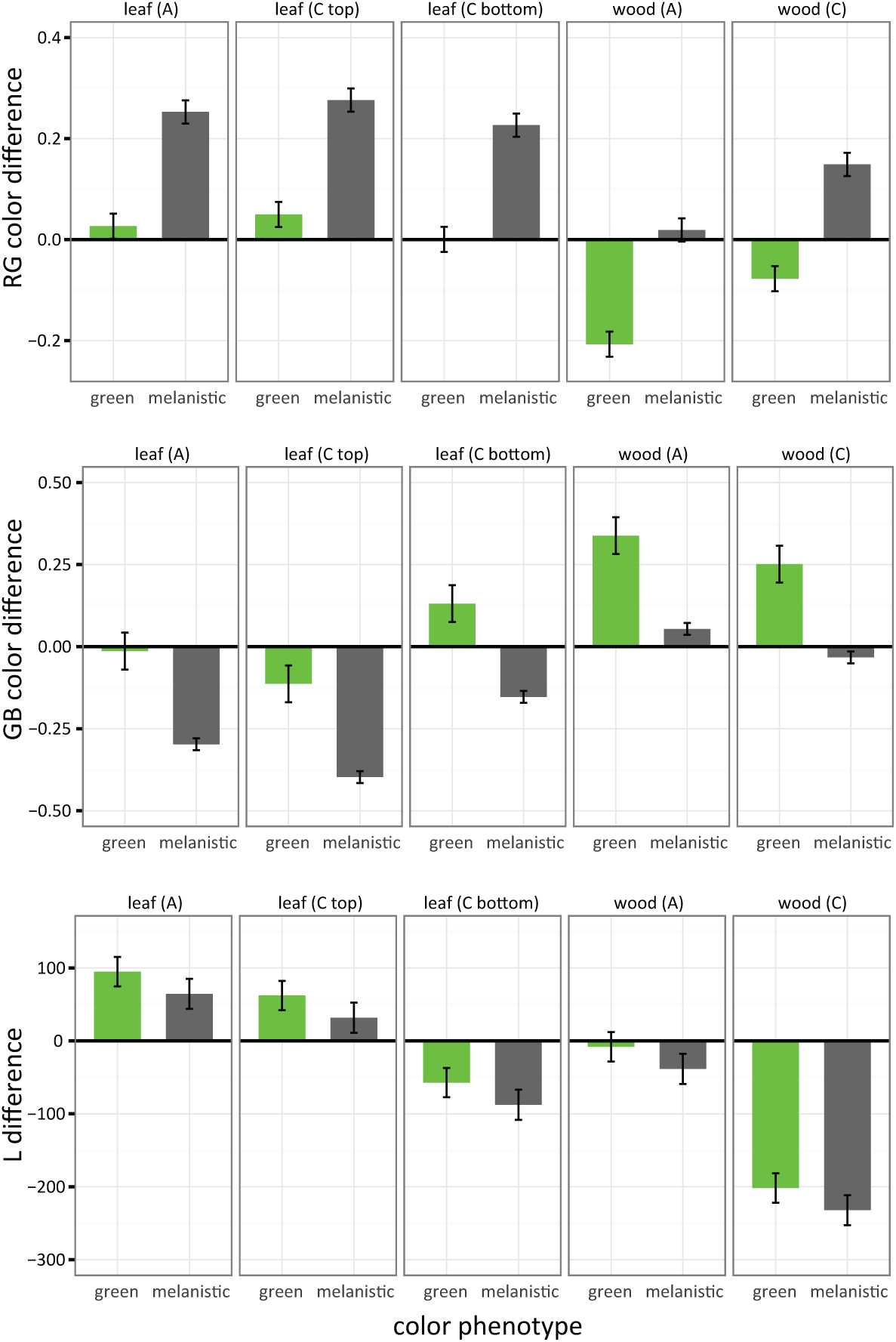
Color matching of *T. podura* color phenotypes to different plant microhabitats. The top row of panels represent RG color differences, the middle panels represent GB color differences, and the bottom row of panels represent L differences. Each row contains 5 panels reflecting the 5 different plant microhabitats we analyzed: *A. fasciculum* leaves (‘leaf (A)’), the top and bottom of *C. leucodermis* leaves (‘leaf (C top)’ and ‘leaf (C bottom)’), *A. fasciculatum* wood (‘wood (A)’), and *C. leucodermis* wood (‘wood (C)’). Bars are mean differences and error bars represent 95% confidence intervals.

### *Genetic architecture of* T. podura *color phenotypes using GWA*

We retained genotypes at 121,435 with minor allele frequency (MAF) greater then 0.01 for GWA mapping analyses. Hyperparameters estimated from BSLMMs indicate that color variation in *T. podura* is controlled by a simple genetic architecture with 97% of phenotypic variation being explained by genotype and 94% of this explained variation being due to only 1 - 4 SNPs with large phenotypic effects (median estimates; Figure 3 for complete posterior distributions). Similar results were obtained for *T. cristinae* with 95% of phenotypic variation in color being explained by genotype and 95% of this explained variation being due to 7 SNPs with large phenotypic effects (median estimates; Figure 3; Comeault *et al.* 2015). Two SNPs in the *T. podura* data set were identified as having a large effect on color phenotypes in > 10% of BSLMM iterations (i.e., PIPs > 0.10). Both of these SNPs map to LG 8 of the *T. cristinae* genome: one at position 10972 of scaffold 1806 (hereafter referred to as “candidate SNP 1”; Figure 4c) and the second at position 349343 of scaffold 284 (“candidate SNP 2”; Figure 4d). The PIP of candidate SNP 1 is 0.295 and the model-averaged estimate of β is 9.92. The PIP of candidate SNP 2 is 0.102 and the model-averaged estimate of β is 4.25.

**Figure 3.**
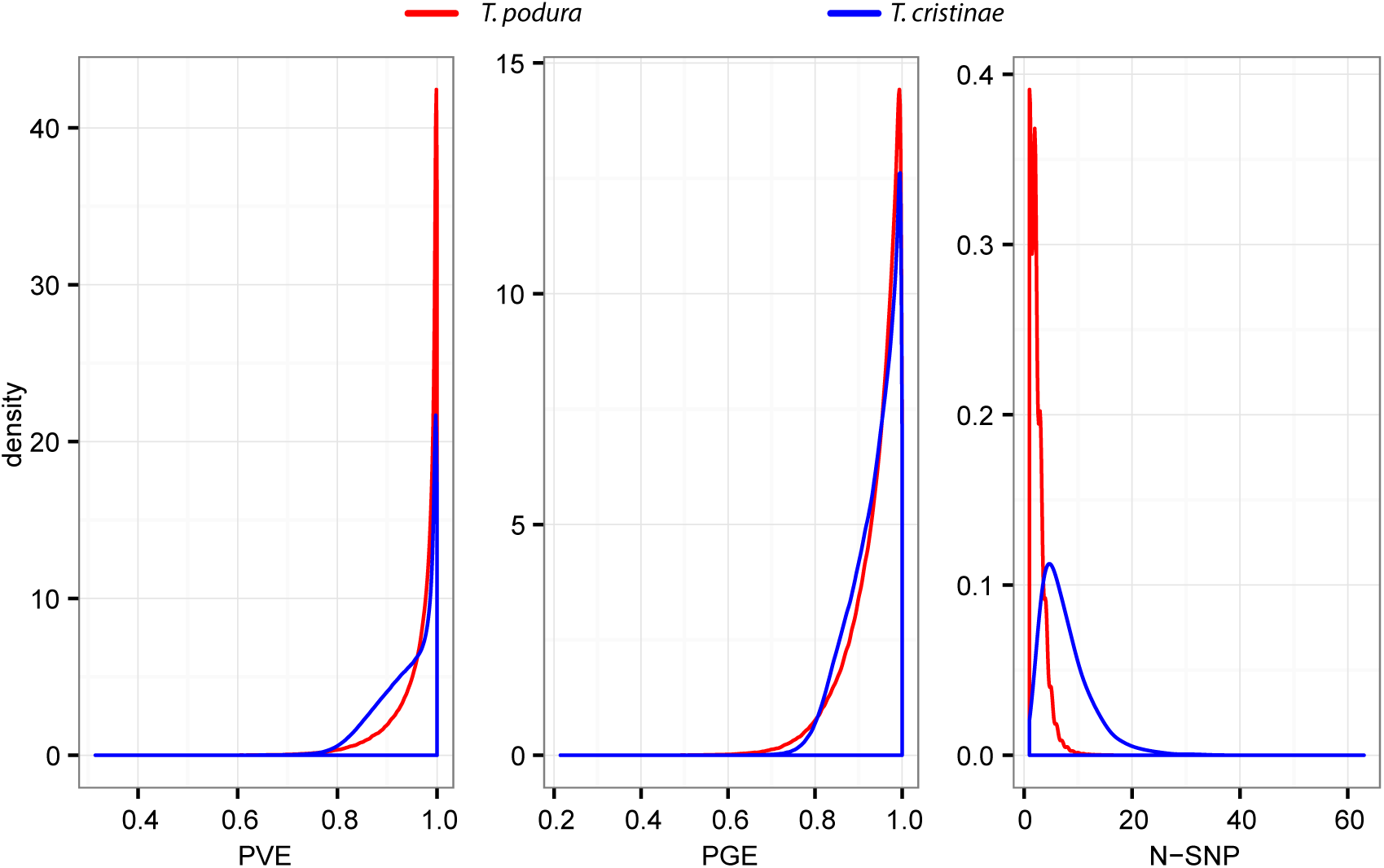
Posterior probability distributions of parameter estimates describing the genetic architecture for color in *T. podura* (red lines) and *T. cristinae* (blue lines). The total amount of phenotypic variation explained by genotype (PVE) and the proportion of that variation that can be explained by SNPs with non-zero affects on phenotypic variation (PGE) are given, along with the number of SNPs in our data set that have non-zero affects on phenotypic variation (N-SNP).

**Figure 4.**
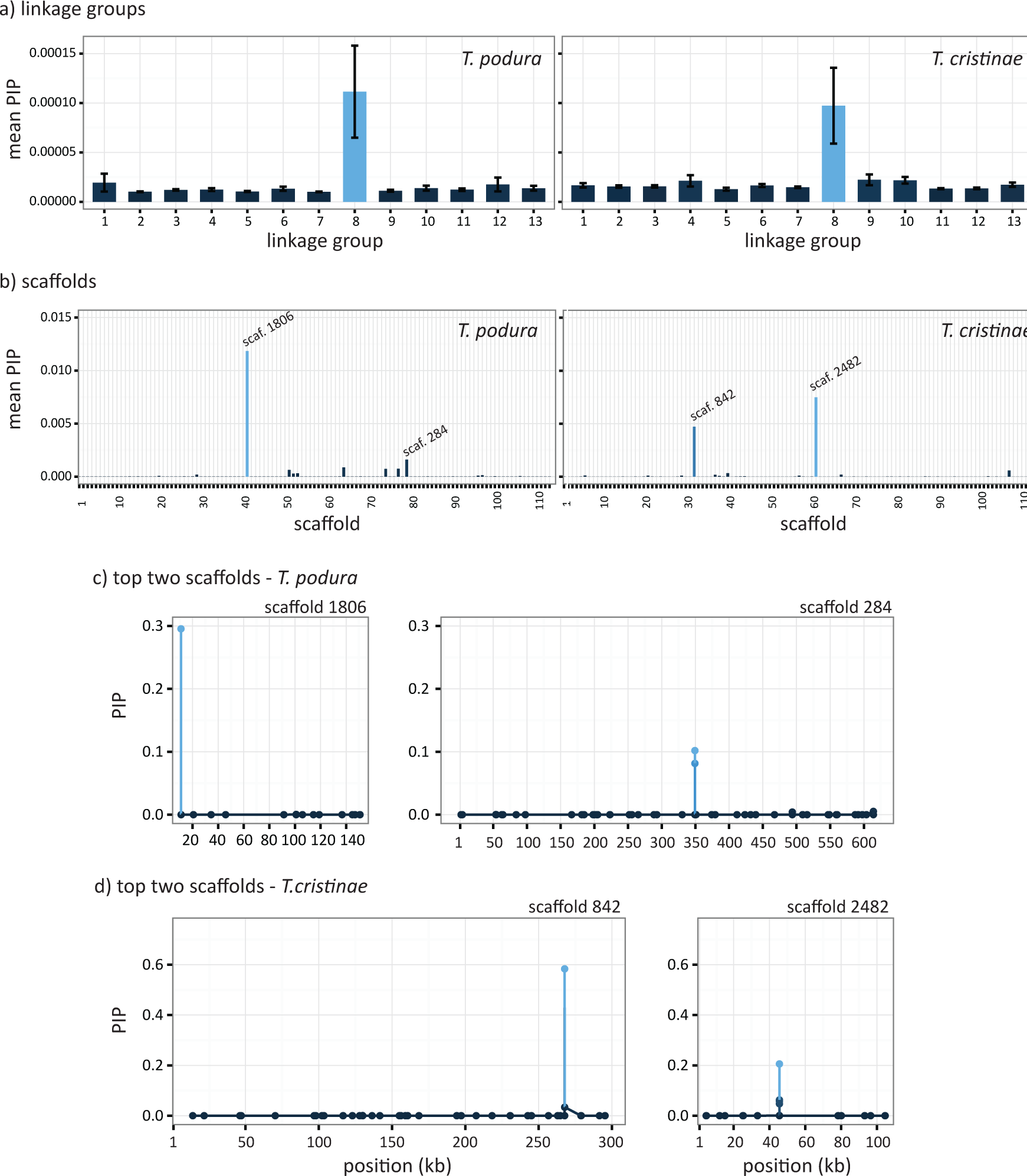
Mapping the genomic location of SNPs associated with color variation in *T. podura* and *T. cristinae*. (a) Mean posterior inclusion probabilities (PIPs) were calculated for SNPs mapping to each of the 13 *T. cristinae* linkage groups (LG) and (b) for each scaffold within LG 8. Scaffolds labels range from 0 to 112 and represent the scaffolds relative position along LG 8. Scaffold (scaf.) number is given for the top two scaffolds for both *T. podura* and *T. cristinae*. Panels c) and d) plot absolute PIP for each SNP mapping to the two scaffolds of LG 8 with the highest mean PIPs in *T. podura* and *T. cristinae*, respectively. Note that *T. podura* candidate SNPs 1 and 2 are represented by the SNPs with the largest PIPs on scaffolds 1806 and 284 (c), respectively.

Further cross-validation analyses revealed effects of this size are unlikely to arise from random associations within our data sets. For example, BSLMM analyses repeated using randomly permuted phenotypic data sets did not recover any SNPs having large effects on phenotypic variation in > 10% of model iterations and confidence intervals for hyperparameter estimates spanned nearly the entire interval [0,1], indicating a strong genetic signal within our observed data (Fig. S3). This strong genetic signal was also confirmed by our ability to accurately predict the phenotype of individuals from genotypic information alone (prediction accuracy = 96.8%; SI; Fig. S4).

We explored whether genomic regions with a large effect on color variation were statistically concentrated on LG 8 by calculating the mean PIP for SNPs at the level of both LGs and scaffolds. Mean PIP differed significantly across the 13 LGs (proportion test; χ^2^ = 21731.33, d.f. = 12, *P* < 0.001). SNPs mapping to LG 8 had the highest mean PIP of all LGs (mean PIP = 0.000111; Figure 4a) and this mean PIP was nearly an order of magnitude greater than the LG with the second largest mean PIP (LG 1; mean PIP = 0.0000194). The two scaffolds with the highest mean PIP were both located on LG 8 and had mean PIPs of 0.0118 and 0.00161 (scaffolds 1806 and 284, respectively; Figure 4b). Notably, these two scaffolds are also the two scaffolds that contain candidate SNPs 1 and 2, respectively.

### Co-localization of regions associated with color in the two species

Further analyses show the co-localization of candidate SNPs in the two species on LG 8 is unlikely to happen by chance. Permutation of PIPs within *T. podura* and *T. cristinae* data sets show that the probability of LG 8 having the highest mean PIP in both species by chance is 0.0067. Within LG 8, the probability that the two candidate scaffolds in *T. podura* and *T. cristinae* are less than or equal to nine and eighteen scaffolds away from each other (the minimum empirical distances we observe between the top two *T. podura* and *T. cristinae* candidate scaffolds; Fig. 4b) by chance was 0.046 ([*P*|distance <= 9]] = 0.156 * [*P*|distance <= 18] = 0.295). These results indicate the same broad genomic region affects color in the two species and raise the possibility the same gene is causally involved.

### Dominance relationships between alleles and linkage disequilibrium analyses

Dominance relationships of alleles at *T. podura* candidate SNPs 1 and 2 show that melanistic alleles are recessive to green alleles: *d*/*a* ratios for the two candidate SNPs identified by BSLMMs are -1 and -0.93, respectively (Figure 5). Genotypes at these two candidate SNPs are also in strong LD (*r*^*2*^ = 0.7917; D’ = 0.9091; Table 2). When compared to genome-wide expectations, estimates of LD between the two candidate SNPs are much greater than mean LD, and tended to be greater than the 95% empirical quantile of LD within each SNP’s respective scaffold, for 1000 SNPs randomly sampled from LG 8, or for 1000 SNPs sampled from across the genome (Table 2).

**Table 1.** Summary of the ecology of the two species of *Timema* stick insects included in this study.

| species | Location | host plants<br>considered here | selection on color<br>phenotypes |
| --- | --- | --- | --- |
| *T. cristinae* | Coastal western<br>Transverse Range,<br>Southern California | *C. spinosus*,<br>*A. fasciculatum* | Balance of multiple<br>sources of selection,<br>often within host species,<br>maintains polymorphism. |
| *T. podura* | San Bernardino, Rosa<br>and San Jacinto<br>Mountains, Central<br>Southern California | *C. leucodermis*,<br>*A. fasciculatum* | Divergent selection<br>acting between host<br>plants. |

**Table 2.** Linkage disequilibrium as calculated as genotypic correlations (*r*^*2*^) and D’ among pairs of loci. Median *r*^*2*^ and D’ is reported for groups of SNPs sampled at different genomic scales (see methods for details; 5% and 95% empirical quantiles are reported for *r*^2^ only). Quantiles were not calculated for the two candidate SNPs as there is only a single pair-wise comparison within this group.

| genomic scale | $r^2$ | D' | 5% quantile | 95% quantile |
| --- | --- | --- | --- | --- |
| candidate SNPs | 0.7917 | 0.9091 | - | - |
| scaffold 1806 | 0.1119 | 0.5505 | 0.0005 | 0.9721 |
| scaffold 284 | 0.0554 | 0.4756 | 0.0002 | 0.2179 |
| LG 8 | 0.0401 | 0.4006 | 0.0002 | 0.1595 |
| genome | 0.0373 | 0.3899 | 0.0002 | 0.1500 |

**Figure 5.**
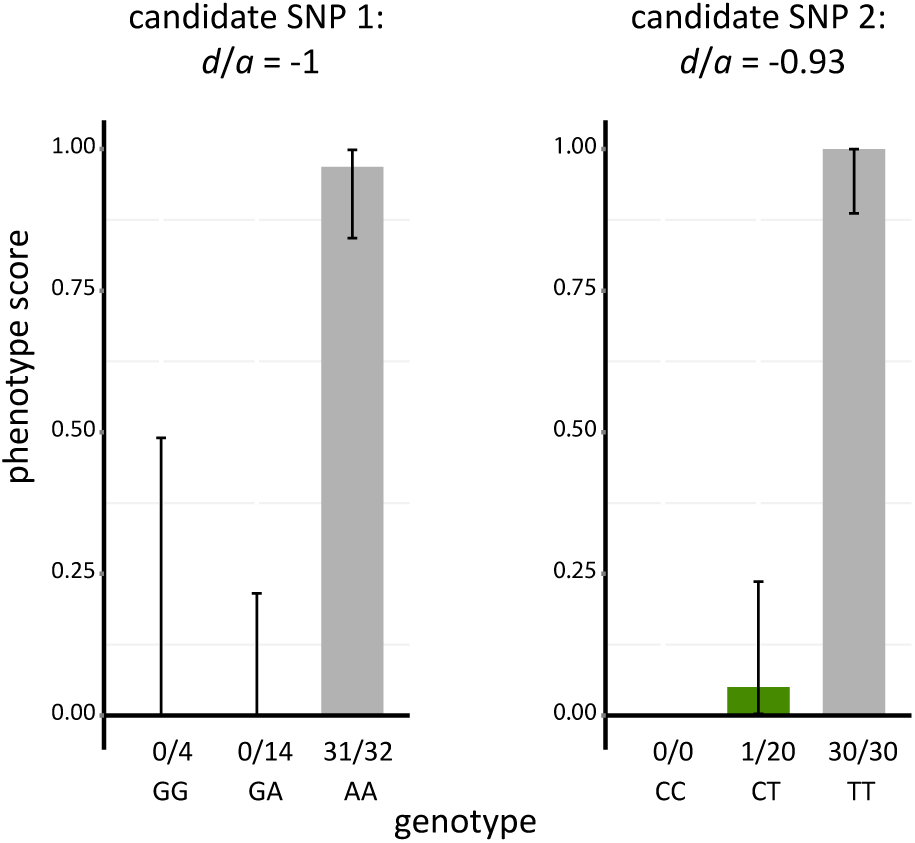
Dominance of alleles at candidate SNPs associated with color variation in *T. podura*. Mean (bars) and 95% binomial confidence intervals (vertical lines; computed using the ‘binconf’ function in R) are shown for each genotype of both candidate SNP 1 and 2. Above each genotype we report the ratio of individuals with that genotype that were melanistic. Green individuals are scored as “0” and melanistic individuals as “1”. See methods for details of how the dominance coefficient (*d*/*a*) was calculated.

### Functional annotation of genomic regions containing candidate SNPs

The two *T. podura* candidate SNPs we identified in this study map to two scaffolds of the *T. cristinae* genome that contain a total of 18 predicted genes with either InterPro or GO annotations (Table S1). However, both candidate SNP 1 and 2 map to intergenic regions and are 13.6 and 4.3 Kb away from the nearest gene, respectively.

Candidate SNPs 1 and 2 do not map to the scaffold of the *T. cristinae* genome containing SNPs most strongly associated with color phenotypes in *T. cristinae* (scaffold 842; Figure 4b and d; Comeault *et al.* 2015). In addition, none of the 39 SNPs that map to this scaffold were located within the putative cysteinyl-tRNA synthetase gene (positions 261250 to 286518) that contained SNPs associated with color variation in *T. cristinae* (Comeault *et al.* 2015). The two *T. podura* SNPs that are closest to this cysteinyl-tRNA synthetase gene are located 1154 bp downstream of its downstream terminus and 8033 bp from its upstream terminus. Median *r*^2^ between the 39 *T. podura* SNPs mapping to scaffold 842 and the two *T. podura* candidate SNPs we identify on scaffolds 1806 and 284 are 0.0204 and 0.0191 (median D’ = 0.3785 and 0.4493) and the maximum pair-wise *r*^2^ is 0.7158 and 0.8502 (maximum D’ = 1.000 and 1.000), respectively. These patterns of LD suggest that the candidate SNPs we identify for *T. podura* tend to be in low LD with SNPs on the candidate scaffold identified in *T. cristinae*. However, some SNPs on this scaffold are in elevated LD with the *T. podura* candidate SNPs and do not preclude the possibility that they are tagging the same functional variation associated with color in *T. cristinae*.

## Discussion

Our results show that similar color phenotypes of *T. podura* and *T. cristinae* largely overlap in two-dimensional color space, with strong divergence between color morphs within species (Figure 1b). In contrast to ‘typical’ parallel or convergent phenotypes (Elmer & Meyer 2011; Losos 2011; Parker *et al.* 2013), the color phenotypes of *T. podura* and *T. cristinae* are not under the same form of selection (Table 1). Therefore, our results highlight how similar phenotypic variation can repeatedly evolve as a result of different forms of selection (Losos 2011). The fact that there are minor differences between green phenotypes of *T. podura* and *T. cristinae* suggests that there are either different alleles at the color locus in the different species (if they indeed share the same locus), ‘modifier loci’, an effect of genomic background, small dietary effects that influence the expression of the green phenotype in these two species, or a combination of these processes.

An important and outstanding question that we were unable to address here is whether color phenotypes of different species of *Timema* evolve via independent *de novo* mutations, ancestral polymorphism, adaptive introgression, or a combination of these mechanisms (Martin & Orgogozo 2013). However, our results suggest that the same genomic regions control color phenotypes in both *T. podura* and *T. cristinae*, and certainly the genetic architecture of these traits shares some similarities between species.

### The genetic architecture of color

We show that color in both *T. podura* and *T. cristinae* is controlled by similar genetic architectures (i.e., one or a few loci on LG 8 with dominance of the green allele; Figure 3). We observed strong LD between the two candidate SNPs identified for *T. podura* (Table 2), suggesting that these two SNPs may be tagging the same functional variant and that color phenotypes of *T. podura* are controlled by a single locus of large-effect. These two SNPs also map to the same linkage group (LG8) as the locus controlling color phenotypes in *T. cristinae* (Comeault *et al.* 2015). We were unable to test whether the same SNPs are associated with color variation in both species because the GBS data set analyzed here did not contain any SNPs within ∼1 Kb of the candidate gene identified in *T. cristinae* (Comeault *et al.* 2015).

Despite not knowing the specific causal variants, some aspects concerning the genetic basis of color are clear. For example, dominance relationships of alleles associated with these color phenotypes are shared between these two species: the green allele is dominant to the melanistic allele (Comeault *et al.* 2015). Moreover, using multi-locus GWA mapping, we have shown that genotype – phenotype associations co-localize to two regions of LG 8 and that this is unlikely to be due to chance. Taken together, these results suggest that the same gene (or group of genes) control color in these two species. Testing this hypothesis will require further fine scale mapping.

Determining the causal mutations controlling phenotypic variation in color in *Timema* would facilitate a better understand the evolutionary history of this variation (e.g., (Colosimo *et al.* 2005; Linnen *et al.* 2009). For example, are green and melanistic phenotypes the result of ancestral polymorphism segregating within populations or are they present due to the recurrent evolution of adaptive color alleles at the same or different loci (Steiner *et al.* 2009)? If this genetic variation represents ancestral polymorphism it would suggest a bias towards the recurrent evolution of the same color phenotypes that affect fitness across different environments. Alternatively, recurrent evolution of the same locus from *de novo* mutation could suggest mutational biases or constraints in the evolution of color in *Timema*. The recent increase in genomic resources available for *Timema* stick insects (Soria-Carrasco *et al.* 2014; Comeault *et al.* 2015) should help to facilitate the discovery of the specific gene or genes underlying these color phenotypes.

### Genetic variation and the response to different selective environments

Dominance relationships at the locus that controls color in the studied species will result in melanistic alleles being hidden from selection in heterozygous individuals. This will likely have two general effects: (1) recessive melanistic alleles will be maintained within populations when they are maladaptive longer than green alleles and (2) dominant green alleles will be able to respond to selection more quickly when at low frequencies in a population compared to melanistic alleles. In *T. podura* the melanistic phenotype, to our knowledge, is fixed within populations living on *Adenostoma* (Sandoval & Nosil 2005). This suggests that there is strong selection acting against the green phenotype on *Adenostom*a, which is supported by past predation experiments (Sandoval & Nosil 2005) and the degree of background matching quantified here (Figure 2), or the green allele has never reached *Adenostoma* populations. By contrast, a combination of either weaker selection against melanistic individuals, the ability of melanistic alleles to hide from selection in heterozygotes, or high rates of gene flow could contribute to the presence of both green and melanistic individuals being found on *C. leucodermis*.

Such influences of genetic architecture on evolution have been shown in *T. cristinae* (Comeault *et al.* 2015) and other systems (Rosenblum *et al.* 2010). For two species of lizard living on the white sands of New Mexico (*Sceloporus undulatus* and *Aspidoscelis inornata*), Rosenblum et al. (2010) showed that dominance relationships of derived *McIr* alleles controlling coloration in these lizard differed: derived ‘white’ alleles were dominant to ‘brown’ alleles in *S. undulatus* but recessive in *A. inornata*. Differences in dominance relationships resulted in different patterns of phenotypic divergence among populations of these lizards adapting to white-sand environments. This example illustrates how understanding the genetic architecture of phenotypic variation can help our understanding of how selection structures genetic and phenotypic variation within and among populations.

### Conclusion

Dissecting the ecological relevance and genetic basis of recurrent evolutionary outcomes (e.g. the evolution of the same color phenotypes in *Timema*) facilitates tests of the factors affecting evolution. Phenotypes that repeatedly evolve among lineages and are controlled by the same genes or genomic regions suggest that certain traits are predisposed to evolve into certain regions of phenotypic space. This pattern could be driven by historical contingencies, pre-existing adaptive variation that segregates within populations (Taylor & McPhail 2000; Blount *et al.* 2008), or genetic and developmental constraints, such as those generated by pleiotropy or epistatic interactions among mutations (Stern & Orgogozo 2009). The data we preset here suggest that color variation in *Timema* stick insects may be highly conserved: *T. cristinae* and *T. podura* are separated by approximately 20 million years of evolution. Ultimately, identifying the specific genes and mutations controlling color phenotypes in *Timema* will help us to better understand the process of local adaptation.

## Acknowledgements

We thank J. Wolf, R. Snook, K.E. Delmore, and members of the D. Matute lab for comments and helpful discussion on previous versions of this manuscript. This research was supported by the European Research Council (Grant R/129639 to P.N.) and the Natural Sciences and Engineering Research Council of Canada (PGS-D3 to AAC). In addition to public repositories, all data are available from the authors upon request.

